# Spectral Discrimination in “Color Blind” Animals via Chromatic Aberration and Pupil Shape

**DOI:** 10.1101/017756

**Authors:** Alexander L. Stubbs, Christopher W. Stubbs

**Author notes:** Both authors contributed equally to the writing of the manuscript.

## Abstract

We present a mechanism by which organisms with only a single photoreceptor, that have a monochromatic view of the world, can achieve color discrimination. The combination of an off-axis pupil and the principle of chromatic aberration (where light of different colors focus at different distances behind a lens) can combine to provide “color-blind” animals with a way to distinguish colors. As a specific example we constructed a computer model of the visual system of cephalopods, (octopus, squid, and cuttlefish) that have a single unfiltered photoreceptor type. Nevertheless, cephalopods dramatically change color both to produce chromatically-matched camouflage and to signal conspecifics. This presents a paradox – an apparent ability to determine color in organisms with a monochromatic visual system – that has been a long-standing puzzle. We demonstrate that chromatic blurring dominates the visual acuity in these animals, and we quantitatively show how chromatic aberration can be exploited, especially through non-axial pupils that are characteristic of cephalopods, to obtain spectral information. This mechanism is consistent with the extensive suite of visual/behavioral and physiological data that have been obtained from cephalopod studies, and resolves the apparent paradox of vivid chromatic behaviors in “color-blind” animals. Moreover, this proposed mechanism has potential applicability in any organisms with limited photoreceptor complements, such as spiders and dolphins.

## Introduction

The only known mechanism of color discrimination in organisms involves determining the spectrum of electromagnetic radiation using differential comparisons between simultaneous neural signals arising from photoreceptor channels with differing spectral acceptances. Color vision using multiple classes of photoreceptors on a two-dimensional retinal surface comes at a cost: reduced signal-to-noise ratio in low-light conditions, and degraded angular resolution in each spectral channel. Thus many lineages that are or were active in low-light conditions have lost spectral channels in order to increase sensitivity (*1*).

Octopus, squid, and cuttlefish (coleoid cephalopods) have long been known to be among the most colorfully-active organisms, vividly changing color to signal conspecifics and to camouflage. In 350 B.C.E. Aristotle remarked (*2*) that the octopus “seeks its prey by so changing its color as to render it like the color of the stones adjacent to it; it does so also when alarmed.”

Cephalopods use their control of skin coloration to become (1) inconspicuous by camouflaging against local backgrounds (Figs. 1, S1 and Movie S1), or (2) highly conspicuous during colorful mating and threat displays (Figs. 1, S2 and Movie S2). Despite this chromatically-active behavior, genetic and physiological studies (*3*–*6*) show that (with one exception) cephalopods lack multiple photoreceptor types. Cephalopods also fail certain behavioral trials (*6*–*10*) designed to test for color vision by opponent spectral channels.

**Figure 1.**
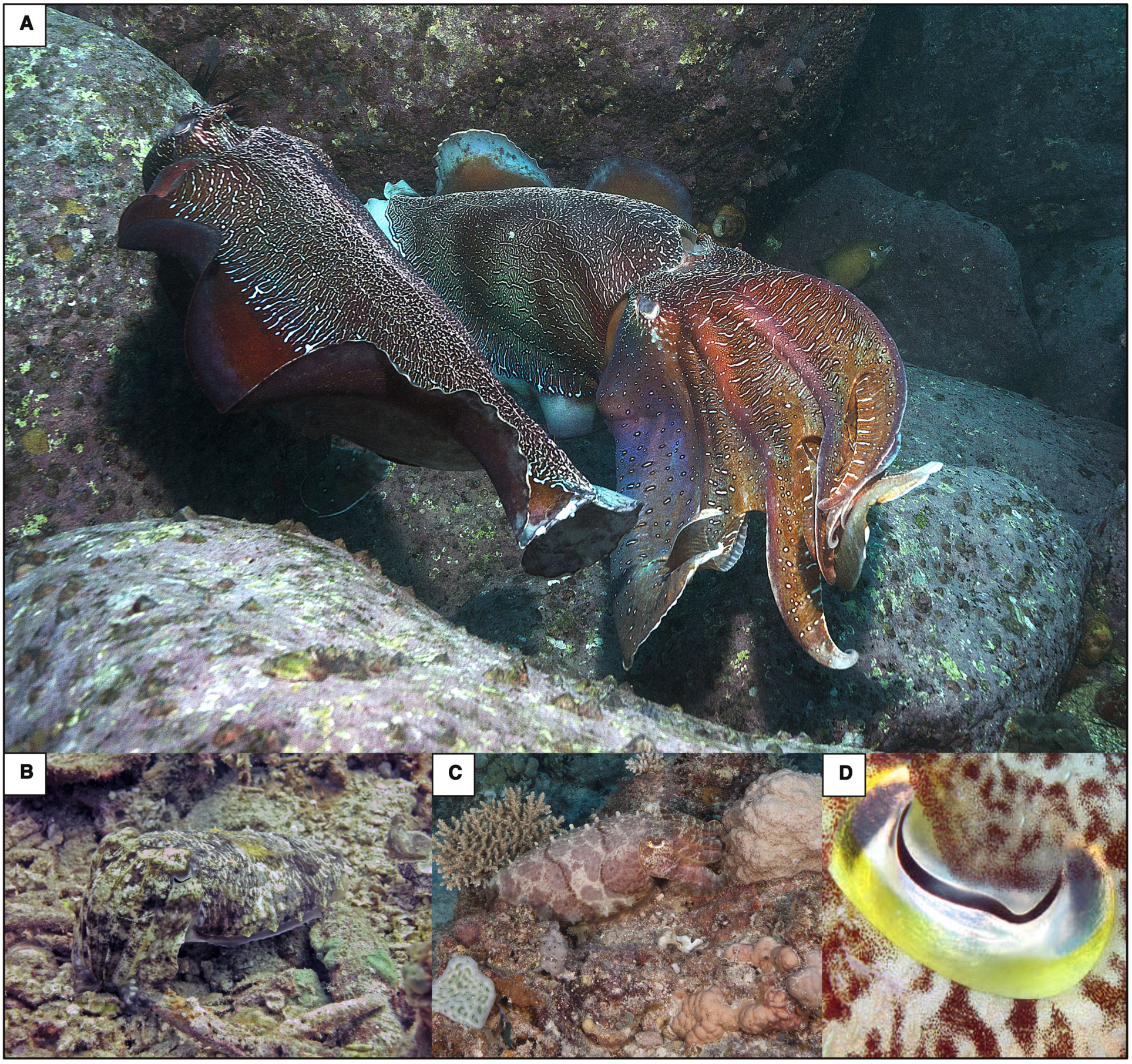
Cephalopod Behavior and Pupil Shapes. Many shallow-water cephalopods produce colorful displays (**A,** Australian giant cuttlefish *Sepia apama*) to conspecifics, and accurately color-match natural environments to camouflage (**B-C**, broadclub cuttlefish *Sepia latimanus*). Their pupil shapes (**D,** *S. latimanus*) maximize chromatic blur. Movies S1-S2 offer additional examples.

This poses two distinct but related paradoxes: (1) how can these animals with a single photoreceptor achieve good background color-matching, and (2) why would they break camouflage to produce risky colorful mating displays (readily visible to predators with color vision) unless this chromatic information were visible to conspecifics and carried some selective advantage?

Previous attempts to reconcile these apparent paradoxes include suggestions that: (1) the animals do not actually match natural background colors (*11*), or (2) multiple photoreceptor types could exist (*12*) in the animal’s skin. Neither of these explanations resolves the puzzle of “color blind camouflage” and researchers remain in search of a mechanism that allows for this ability (*6*, *13*–*15*). We are unaware of a proposal for how natural selection would drive the evolution and maintenance of colorful intraspecific displays in these soft-bodied mollusks if this information were not available to the animals themselves.

We assert in this paper that under certain conditions organisms can determine the spectral composition of objects, even with a single photoreceptor type. Through computational modeling we show how this mechanism provides spectral information using an important relationship: the position of sharpest focus depends upon the object’s spectral peak. Mapping out contrast vs. focal setting (accommodation) amounts to obtaining a coarse spectrum of objects in the field of view, much as a digital camera attains best-focus by maximizing contrast vs. focal length.

**Contradictory evidence: chromatic behavior but a single opsin.** The extent of color matching in cephalopods remains somewhat controversial in some circles, but we assert that shallow-water cephalopods often match the coloration of natural backgrounds (Figs. 1, S1 and Movie S1), and we encourage readers to examine the Supplementary Materials footage (Movie S1) of cuttlefish and octopus camouflage in their natural habitat and reach their own conclusions. Some have claimed (*11*) that these organisms simply match the brightness and spatial scale of patterns in their environment, tricking the human visual system without actually requiring a color match. Numerous studies (*13*, *16*–*20*) show, however, that cuttlefish and octopus actively vary their spectral reflectance in response to background color rather than simply modulating their luminance.

Kühn conducted (*20*) a series of behavioral experiments comparing the octopus and cuttlefish camouflage response when placed on a series of greyscale and colored substrates. His data show statistically significant evidence that these organisms expand their long-wavelength-reflecting chromatophores when on red-yellow backgrounds, but primarily expand black chromatophores when on greyscale backgrounds. He concluded that these organisms must have the ability to discriminate spectral content.

Contemporary laboratory and field observations (*16*–*19*) show octopus and cuttlefish produce high-fidelity color matches to natural backgrounds (Fig. 1). The most definitive recent evidence for color camouflage matching in a laboratory setting used (*13*) a hyperspectral imager in conjunction with spectral angle mapping to show cuttlefish varied their spectral reflectance (chromatic properties) to maintain excellent spectral matches to a diversity of natural backgrounds, and interestingly maintained poorer matches in brightness (luminance). These studies (13, 16–19) corroborate Kühn’s earlier result: cephalopods actively vary their spectral reflectance via active control over their chromatophores in response to natural backgrounds, rather than simply varying their luminance.

Rhodopsin is expressed (*12*, *21*) in both octopus and cuttlefish skin and, as in many other mollusks, this skin has the ability to detect the intensity of light. The skin’s spectral response was shown (*21*) to be nearly identical to that of the eye. There is no evidence or proposed mechanism for how this single opsin in the skin would contribute to spectral discrimination and allow for spectrally-matched camouflage. Additionally, absent a focusing element, detectors on the skin act as wide-angle non-imaging light sensors and cannot provide useful information regarding background coloration or signals produced by conspecifics.

**Chromatic blur dominates image quality budget.** A variety of factors determine the blurring of the image formed on the retina, including diffraction through the pupil, aberrations in the optical system, and retinal limitations (*1*, *22*–*29*). These terms comprise the image quality budget and determine the sharpest image that can be formed. The eyes of *Octopus australis* are particularly well-studied (*22*) and we used data from this species as a proxy for other shallow-water cephalopod eyes to make quantitative assessments of the image quality budget. The *O. australis* lenses have two properties shared by all other studied cephalopods: (1) they are remarkably well-corrected for spherical aberration, and (2) the index of refraction varies with wavelength, inducing chromatic aberration. This chromatic aberration is uncorrected, and is found in all studied (*23*–*24*) cephalopod lenses. In some other animals radial multi-focal zones do produce a partial chromatic correction (*25*).

Wavelength-dependence of the index of refraction induces (*22*) chromatic blurring, since different wavelengths have different focal lengths. This effect dominates the image quality budget (Table S1). The extent of chromatic blurring depends on both chromatic focal shift and the angle at which rays strike the optical axis (Fig. 2). This angle depends on the ray’s height *h*; off-axis pupil area determines the extent of chromatic blur. Even though the single opsin restricts the range of wavelengths detected, our analysis demonstrates that when integrated over the wavelength response, chromatic blurring dominates image quality except for small, on-axis pupils or when the lens diameter is so small that the granularity of the photoreceptors dominates. A monochromatic point source generates a scaled image of the pupil, with both size and parity determined by the amount of defocus (Fig. 2, Figs. S4-S6).

**Figure 2.**
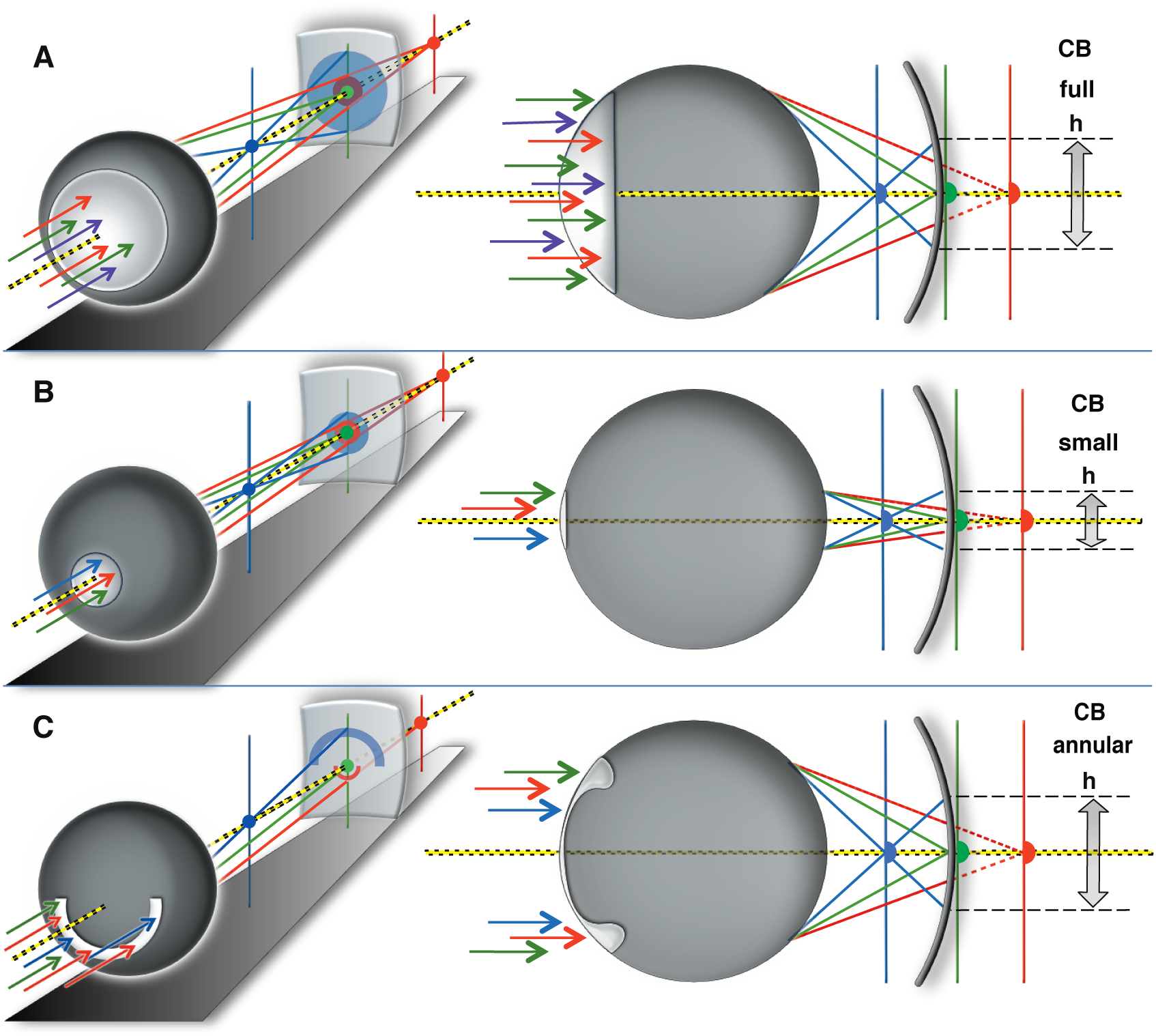
Chromatic Blur and Pupil Geometry. The full (**A**) and annular (**C**) aperture pupils produce more chromatic blurring (CB) than the small on-axis (**B**) pupil because they transmit rays with a larger ray height *h*. Vertical lines show best-focus positions for blue, green, and red light.

**Range, color, and best-focus.** Organisms routinely determine the best-focus for objects of interest in their visual field by varying focal length and comparing relative image quality. This can be used as an accurate range-finding mechanism, as shown in chameleons (*30*) and jumping spiders (*31*). If chromatic blur dominates the image quality budget, there is an inter-relationship between range, color, and best-focus (Fig. 3). For example jumping spiders misjudge (*31*) distance depending on the illumination spectrum. Differential image blurring has been proposed (*32*) as a range-finding mechanism for squid, but chromatic aberration (not considered in Chung & Marshall (*32*)) drives a strong relationship between spectrum, range, and best-focus. Even in this narrowband system chromatic aberration can compromise the determination of range based on best-focus values.

**Figure 3.**
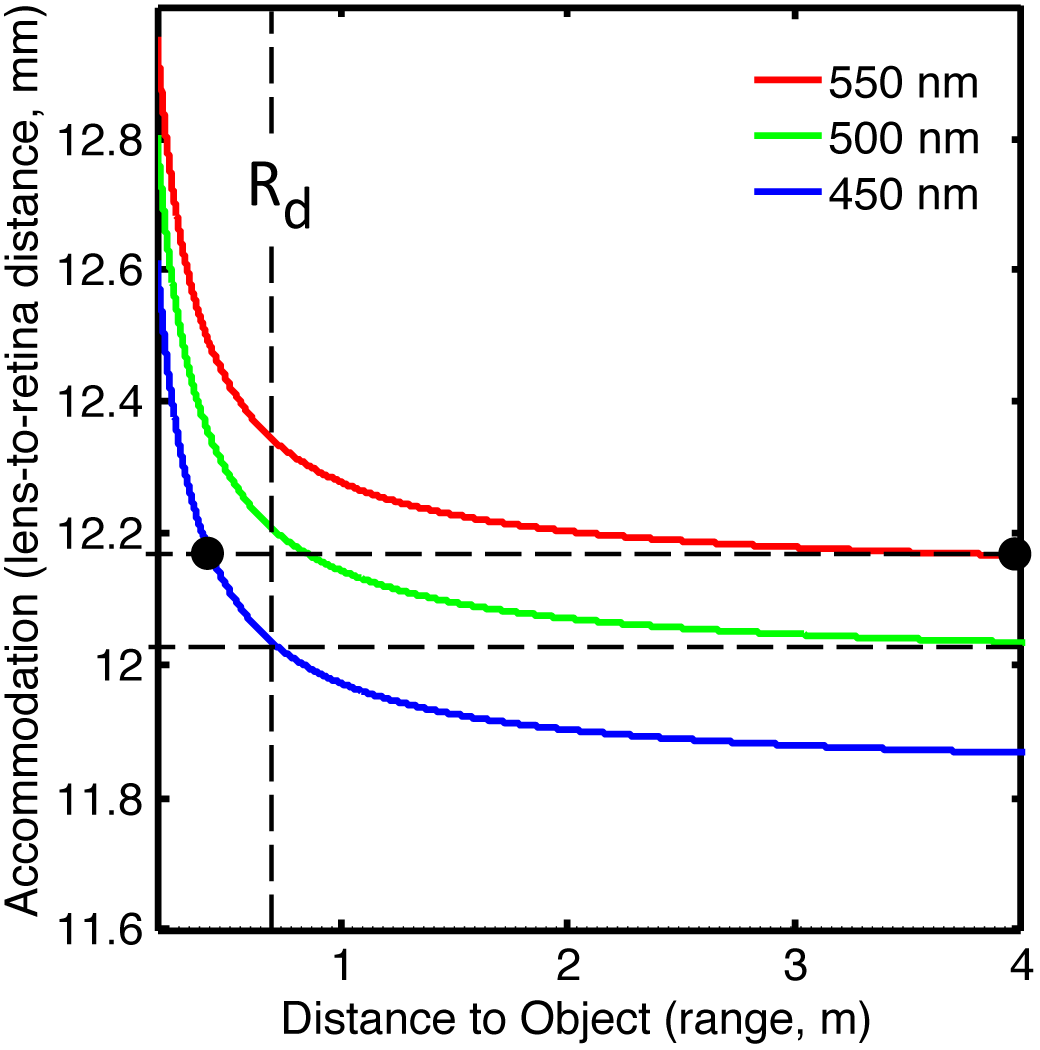
Range-color-focus relationship for a 10-mm diameter cephalopod lens. Colored lines show the accommodation vs. range relationship at the 500-nm opsin peak sensitivity and 450-nm and 550-nm 50% sensitivity points. For objects more distant than R_d_, there is an unambiguous relationship between wavelength and the accommodation setting that makes the sharpest image.

Coleoid cephalopods can use (*33*) binocular vision to judge distance. This can help resolve the color-range ambiguity. The combination of a determination of best-focus and a rough determination of range allows for spectral discrimination.

The spherical lens system we modeled obeys a modified lensmaker’s equation. The image distance *I*, is a function of the wavelength-dependent focal length *f(λ)* and object distance *O*, with

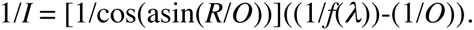

We used this expression and the measured (*22*) chromatic aberration for *O. australis* to compute the image distance *I* needed to achieve a focused image as a function of both object distance and wavelength at λ=450, 500, and 550 nm. These wavelengths correspond to the opsin peak and the 50% sensitivity points. Figure 3 shows the relationship between object distance (range) and the lens-to-retina spacing (accommodation) for these chosen wavelengths. Although there is color-range ambiguity for nearby objects, range-independent spectral discrimination (defined here as discrimination between the opsin peak wavelength and the two 50% sensitivity points) can be achieved for objects at distances beyond R_d_=(d/(10mm))(0.75m), for lens diameter d. In this geometrical optics regime, the disambiguation range R_d_ scales with lens diameter.

Beyond R_d_, best-focus depends only on spectrum and is independent of range. A scan through focus amounts to a spectral scan of the scene. The animal can determine the object’s color by finding the focal setting that produces the sharpest image, regardless of range. This best-focus determination can be achieved either by displacing the lens (*34*) relative to the retina (accommodation), or by imaging the object across regions of the retinal surface with different effective focal lengths.

What occurs when objects are closer than Rd? The focal spacing creating a crisp focus of a 450-nm light source at 0.2-meters also creates a sharp image of a 550-nm light source more than 4 meters away (Fig. 3). An independent distance determination, even if coarse, can break this ambiguity. Studies of range determination in cuttlefish show they utilize multiple methods for precisely establishing distances (*33*). Both cuttlefish and squid rely on this ability to accurately project their tentacles and capture food. This ranging ability can break the range-color degeneracy and allow them to use image sharpness to obtain spectral information for R<R_d_.

This mechanism for spectral discrimination is computationally more intensive than a differential comparison of photoreceptor outputs in opponency. We believe this may be one factor contributing to the exceptionally large (*11*) optic lobes found in coleoid cephalopods.

**Modeling spectral sensitivity from image sharpness.** We computed the relationship between image sharpness, accommodation, and spectral content. We created test patterns with different spectral characteristics and simulated the images they would form on the single-opsin retina of *O. australis*, for different accommodation values. The test patterns (Fig. 4A-E) are generated with the reflectance spectra (*35*) of blue and yellow Australian reef fish. The side length of each pixel in the test patterns is equal to the 5-micron rhabdome diameter; our test images incorporate the sampling granularity inherent in this detector system.

**Figure 4.**
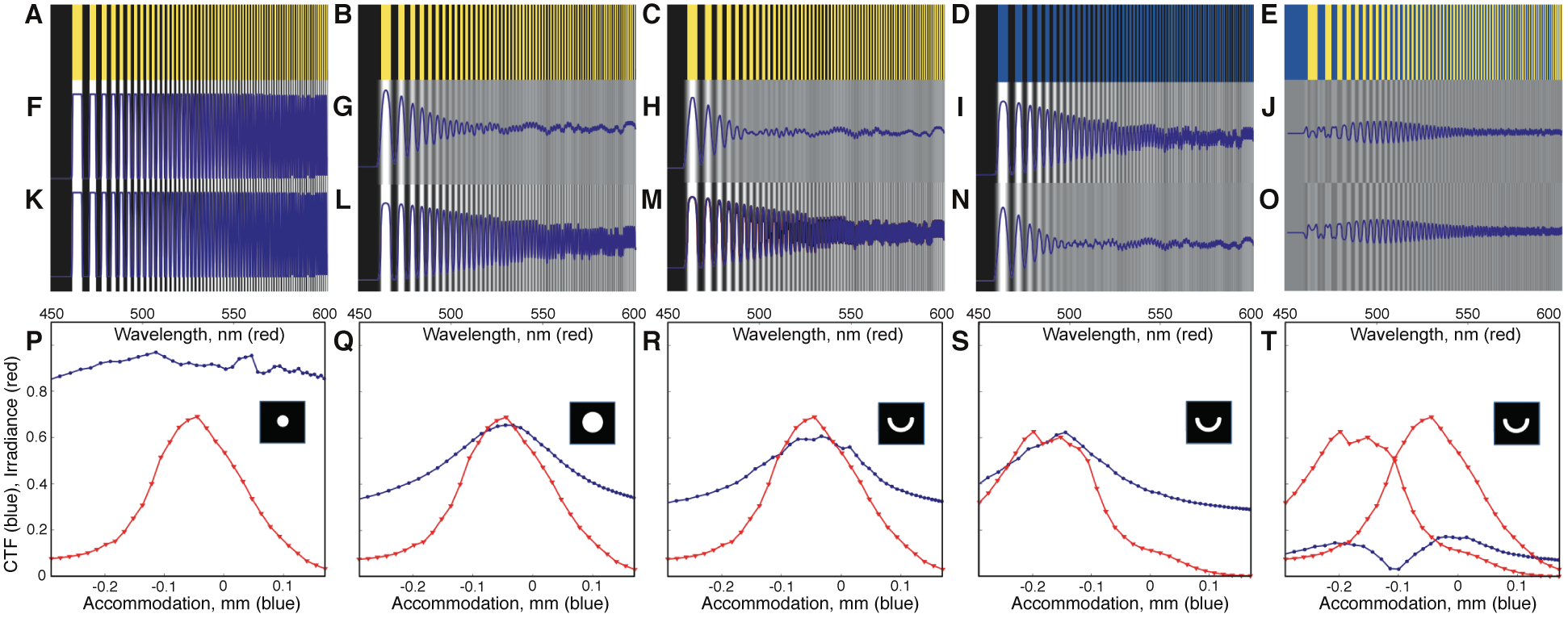
Chromatic blur simulations for semi-annular pupil. Test patterns (**A-C** in black-yellow, **D** in black-blue, and **E** in blue-yellow) used to simulate chromatic blur vs. accommodation. Examples of detected intensity variations and contrast at best-focused wavelengths of 470-nm (**F-J**), 550-nm (**K-O**). CTF is extracted from line plots of intensity (blue traces in **F-O**). CTF (blue, **P-T)** vs. accommodation (lower x-scale) tracks the spectrum of detected photons (red, **P-T**) vs. wavelength (upper x-scale), with a spectral resolution that depends on pupil shape. The pupil-dependence of spectral resolution (width of blue CTF traces) is shown for small (d=1mm, **P**) and full (d=8mm, **Q**) on-axis pupils, and for the semi-annular (6mm<d<6.66mm) pupil (**R-T**). The CTF peak tracks the spectrum (**R-T**) for the semi-annular pupil. The flat CTF vs. accommodation (**T**) obtained from the line plots of intensity (**J, O**) for the blue-yellow test pattern (**E**) precludes spectral discrimination for this case.

Movie S3 is an animation that shows how the contrast of the image depends on focal setting (*i.e.* accommodation). Maximum contrast is obtained when the lens brings light at the peak of the detected photon spectrum into best-focus. Two representative simulated blurred images are shown in Figure 4 for each of the five test patterns, for focal settings that bring 470-nm and 550-nm light into focus on the retina. The amount of chromatic blurring is evident from their corresponding intensity line cuts, shown as blue superimposed lines in the respective figures. We mapped image sharpness vs. accommodation setting over the full range of wavelengths. The spectral peak of the light detected from an object can be inferred from the accommodation setting where the image is best-focused on the retina.

We computed a Contrast Transfer Function (CTF, Fig. 4) metric to map out image contrast as a function of accommodation, pupil shape, and the spectral content of the test image. The CTF vs. accommodation for a given test pattern tracks the underlying spectrum but with a spectral resolution that depends on intensity contrast as well as pupil shape. Figure 4P-T show the spectral content of the test patterns (red lines), and the computed image-sharpness vs. accommodation (blue lines).

Under our model the determination of spectral information is reliant on fine-scale intensity variations (edges, shadows, texture, etc.). This imposes limitations. Cephalopods would be unable to determine the spectral content of a flat-field of uniform color. They would similarly be unable to determine spectral information from abutting regions of comparable apparent intensity, differing only in spectral content (Fig. 4E, J, O, T). This can account for results obtained in laboratory behavior tests for color vision (Table S2).

Natural environments rich in shadows and structure serve as focusing aids. Spectra measured (*35*) in marine environments often provide the spectral structure needed for this mechanism. Intraspecific displays of these organisms (Figs. 1, S2 and Movie S2) typically exhibit adjacent fine-scale black and colored regions, facilitating best-focus determination. We believe this is another adaptation that favors our model.

**Chromatic blurring favored over visual acuity.** While ambient light levels influence optimal pupil area, pupil *shape* determines the extent of chromatic blurring (Fig. 2). Chromatic blur dominates the cephalopod image quality in low-light conditions, with a fully-dilated pupil (Table S1). The off-axis slit and semi-annular pupils used in high-light conditions preserve this spectral discrimination mechanism across a wide dynamic range of illumination. The semi-annular pupil shape (Fig. 1), common in both cuttlefish and shallow-water squids, maximizes the off-axis distance of optical rays from objects in the horizontal plane around the animal. The horizontal slit pupil of shallow-water octopus (Fig. S2F) species intercepts a similar ray bundle from when imaging the bottom, acting as an arc-like pupil for images formed on the upper portion of the retina that has an enhanced density of photoreceptors (*36*).

We computed the pupil dependence of CTF vs. accommodation, and the corresponding spectral resolution, for three pupil shapes using the yellow-black test pattern (Fig. 4A-C). The small on-axis pupil (Fig. 4P) has minimal chromatic blur and maintains a crisp CTF across the range of accommodation settings, maximizing visual acuity but with degraded spectral sensitivity. Full (Fig. 4Q) and semi-annular pupils (Fig. 4R) more realistically represent cephalopod pupils under low-light and high-light conditions respectively, and have virtually identical accommodation-dependent chromatic blur, and correspondingly higher spectral resolution than the small pupil.

We propose that natural selection might favor the maintenance of spectral discrimination over image acuity in these animals.

**Consistency with experiments.** Despite earlier behavioral results indicating color discrimination (*8*, *20*), two lines of evidence drove (*6*) the prevailing view that nearly all cephalopods are color blind. First, only one photoreceptor type exists (*3*–*5*) in the retina of shallow-water cephalopods. Our mechanism for spectral discrimination requires only one receptor type. Second, some behavioral experiments (*6*–*10*) designed to test for color vision in cephalopods produced negative results by using standard tests of color vision to evaluate the animal’s ability to distinguish between two or more adjacent colors of equal brightness. This is an inappropriate test for our model (Fig. 4R). Tests using rapidly vibrating (*7*–*8*) color cues are also inappropriate. While these are effective tests for conventional color vision, they would fail to detect spectral discrimination under our model because it is difficult to measure differential contrast on vibrating objects. These results corroborate the morphological and genetic evidence: any ability in these organisms for spectral discrimination is not enabled by spectrally-diverse photoreceptor types. Table S2 reviews cephalopod behavioral experiments investigating color vision and their consistency with our proposed mechanism.

**Other organisms.** Our proposed mechanism has potential applicability in other species with a limited number of photoreceptor types and low *f*-number visual systems. Some dolphin species utilize (*37*) an annular pupil and a similar (*38*) radial-gradient index of refraction lens uncorrected for chromatic aberration. They display evidence for behavioral color discrimination (*39*) in spectral regimes where their visual system would have difficulty (*40*) encoding color by opponent channels. More generally, a large number of organisms that are active both diurnally and nocturnally possess (*41*) an annular pupil, and we wonder if these organisms could also benefit from color discrimination by our proposed mechanism.

Camera-style eyes with a fast lens and a retinal surface with little or no spectral discrimination arose independently across multiple lineages. Even in the single-opsin case, chromatic effects can dominate image blur, albeit by an amount dependent on pupil shape.

Spider primary eyes use a low ƒ-number optical system and thus induce high chromatic blurring, maintained by a “boomerang” shaped (42) pupil that maintains this chromatically induced defocus. Additionally, all studied spiders image their environment with only two functional opsins (UV and green peak sensitivities) (*31*), though a recent study (*42*) shows that one genus of jumping spider may use retinal filtering to obtain color vision by opponency on a tiny portion of one focal plane. Spiders also use an imaging system that maintains (*42*) off-axis rays in high-light conditions (as in the cephalopod annular pupil) and simultaneously (*43*) image across multiple axially-displaced focal planes. Jumping spiders use (*31*) image defocus across these focal planes to judge distance, but (as in cephalopods) this can be confounded by chromatic aberration due to color-range ambiguity (Fig. 3). Simultaneous imaging across multiple focal planes provides chromatic information by comparing the relative defocus of objects across these focal planes. Many spider species image across 3–4 focal planes, more than would be required if the tiering were simply to correct for chromatic aberration in each of the two photoreceptor channels. By determining the retinal tier with the best-focused image spiders should be able to make spectral inferences without requiring multiple photoreceptor classes acting in opponency in a given spectral region.

Some jumping spider genera (such as peacock jumping spiders which lack retinal filtering pigments) exhibit displays in various shades of green and red. This is hard to account for in an organism that would conventionally be red-green color blind. While under natural sunlight, some jumping spiders exhibit a preference for red-colored mates (*44*) and crab spiders show an ability to background-match (*45*). However, these behaviors disappear under fluorescent lighting. Fluorescent lighting in these experiments created a series of line emissions that approximate δ functions and dominate the reflected spectrum from objects in the visual field, and this would make spectral inferences by chromatic defocus imaging difficult. By simultaneously comparing image quality across multiple offset focal planes they might be able to obtain more spectral information than by two photoreceptors working in opponency, and indeed tiered retinas found (*1*) in spiders and many deep-sea fish likely represent the optimal morphology for spectral discrimination using our proposed mechanism.

We believe that in many organisms chromatically-induced blurring may offer an additional source of spectral information, forcing us to rethink what it means to be a “color blind” animal.

## Acknowledgments

AL.S. thanks University of California at Berkeley and the Museum of Vertebrate Zoology for their support, and is grateful to Professor J. McGuire for extensive opportunities, mentoring, and comments on this manuscript, and to R. Caldwell and M. Banks for helpful comments. C.W.S. acknowledges the support of Harvard University. S. Johnsen provided insightful comments for which we are very grateful. We thank N.O.S.B. and the Packard Foundation for their support of science, J. Schoeneberg for translation assistance, and E.C. Gregory for editorial support. Photographs in Fig. 1 were provided by (A) K. Stiefel, (B, D) L. Sawitri, and (C) K. Marks. Photographs in Fig. S1 were provided by (A) K. Marks, (B-D) L. Sawitri, (E) K. Stiefel, and (F) R. Caldwell. Photographs in Fig. S2 were provided by (A, C) Oceanwide Images, (B) R. Tan (Wild Singapore), and (D) R. Ling. We thank R. Hanlon for permission to include Movie S1, and J. Aguilera for making Movie S2 similarly available. The MATLAB routines used for computing the results shown here are available on github (link to be made public upon publication).

## Inventory of Supplementary Materials

### Methods and Materials

Chromatic Aberration Computation
Illumination
Reflectance Spectra and Test Images
Point Spread Function (PSF)
Contrast Transfer Function (CTF) Analysis
Image Quality Budget

Photoreceptor size
Retinal displacement
Residual spherical aberration
Chromatic Aberration
Diffraction
Other Achromatic Aberrations
Consistency with previous behavioral experiments

Figures
Figure S1. Color matching and pupil shape in cephalopods
Figure S2. Examples of highly colorful signaling in cephalopods
Figure S3. Spectra used in simulations showing the wavelength dependence (in nm) of multiple quantities of interest
Figure S4. Chromatic PSF for 400-nm best-focus, semi-annular pupil
Figure S5. Chromatic PSF for 700-nm best-focus, semi-annular pupil
Figure S6. Chromatic PSF for 700-nm best-focus, full pupil
Figure S7. Encircled energy vs. radius for various accommodation settings

Tables
Table S1. Cephalopod retinal image quality budget
Table S2. Prior behavioral Cephalopod experiments testing for color vision

Movies
Movie Caption S1. *Octopus vulgaris* demonstrating dynamic background color matching in the wild
Movie Caption S2. Cuttlefish *Sepia latimanus* demonstrating chromatic signaling to conspecifics
Movie Caption S3. Contrast dependence on focal setting

## Materials and Methods

### Chromatic Aberration Computation

Ideally, a set of monochromatic measurements of the Point Spread Function (PSF) produced by a cephalopod lens, for different pupil sizes and lens-to-retina spacings, would establish an empirical determination of the chromatic blur seen by these creatures. We are unaware of an appropriate comprehensive data set, so we have used the available laboratory measurements to produce a computer model of the chromatic properties for a representative cephalopod. Since the primary eye design features (complex pupil shape, spherical gradient-index lens, and single-opsin retina) are common across most cephalopods, we will use this model as representative of this class of animals.

Using measured (*22*) optical properties of *Octopus australis* we performed a simulation by constructing a hyperspectral image cube (at 5 microns/pixel in the spatial directions, corresponding to a typical cephalopod rhabdome diameter (*11*, *46*), and 200 planes spanning 450<λ<650 nm in the spectral direction at Δλ=1 nm). We modeled an *f*/1.2 spherical lens with a 10-mm diameter, but our computed chromatic blurring results are independent of this choice of length scale. For each lens-to-retina focal distance, which brings a single wavelength into crisp focus, we computed the pupil-dependent chromatic image blur at the other wavelengths. We summed up the blurred image cube along the wavelength direction, weighted by the product of the seawater-filtered solar photon illumination, the reflectance spectrum, and the opsin response curve (Fig. S3), to arrive at a final simulated chromatically blurred image on the retina. This procedure was repeated for three different pupil shapes for a sequence of accommodation values.

Chromatic blurring (Fig. 4) was computed with a MATLAB code, chromatic.m, adapted from a program initially written by C.W.S. to investigate the out-of-focus properties of the Large Synoptic Survey Telescope (LSST). The PSF and encircled energy diagrams (Figs. S4-S7) shown below were computed with a related program, PSF.m.

The detected light intensity I(i, j) in each pixel (i, j) of the simulated retinal image is given by

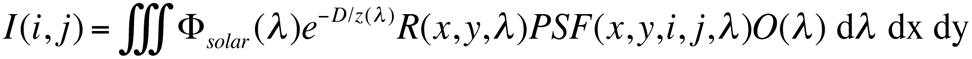

where F_solar_(λ) is the solar photon irradiance spectrum, the exponential term accounts for the reduction in down-welling photon flux at a water depth D with an attenuation length z(λ), R(x, y, λ) is the spectral reflectance of the portion of the scene at a location x, y, PSF(x, y, i, jλ) is the wavelength-dependent PSF of the visual system at pixel (i, j) for light arriving from location (x, y) in the scene, and O(λ) is the sensitivity function of the photoreceptor opsin. The limits of integration span the spatial extent of the scene within the field of view and the spectral range of interest, in this case band-limited by the opsin response O(λ).

### Illumination

We used a ground-level spectrum of solar irradiance from http://www.pvlighthouse.com.au/resources/optics/spectrum%20library/spectrum%20library.aspx, multiplied by λ to convert to relative photon irradiance. To account for illumination attenuation at a 3-m depth D, we used the Pacific seawater optical attenuation length data from Morel & Maritonrena (*47*). We used the attenuation lengths appropriate for 0–20 meter depths, which correspond to a 0.043 mg m-3 chlorophyll density. The resulting photon irradiance spectrum is presented in Fig. S3A.

### Reflectance Spectra and Test Images

Each pixel in the simulated test image was assigned a reflectance spectrum that was a weighted superposition of template reflectance spectra. To simulate the colors encountered in marine settings we use measured (*35*) reflectance spectra of blue and yellow tropical fish.

The reflectance spectra are shown in Fig. S3C. We produced a variety of dual-color bar charts (shown in Fig. 4A-E) with varying spatial frequency in order to assess the acuity of the resulting images. The spatial scale of these test images was set to 5 microns/pixel, which corresponds to the typical diameter of the photosensitive structures (rhabdomes) that pave the retinal surface of cephalopods. We produced bar chart test patterns with 112-pixel to 2-pixel bright bar widths. At our sampling of 5 microns/pixel this corresponds to 165-micron and 10-micron widths, respectively, on the retina.

### Point Spread Function (PSF)

The imaging properties of the cephalopod visual system are determined by the combination of the pupil shape, the refractive properties and configuration of the optical components, and the shape and location of the retinal surface. Rays first propagate through the pupil, which determines both the collecting area and the off-axis distances *h* of the rays that are imaged onto the retinal surface. The cephalopod lens is well-approximated by a sphere with a radial gradient in the index of refraction that produces a remarkably effective correction for spherical aberration (*22*). The wavelength-dependence of the index of refraction does, however, produce chromatic aberration. Different wavelengths have different effective focal lengths. The blur induced by this depends on the angle at which the rays intersect the optical axis, which in turn scales with the ray’s distance off-axis. Pupils that transmit a large proportion of off-axis rays produce more chromatic blurring than pupils that are predominantly on-axis.

When chromatically out-of-focus rays emanating from a point source at infinity of a given wavelength intersect the retina, they produce a PSF that is a scaled image of the *pupil* (Fig. 2). The resulting focal plane image at each wavelength is a convolution of this wavelength-dependent PSF with the test pattern image. We computed this convolution for a discrete set of wavelengths, and summed the resulting hyperspectral synthetic image along the spectral direction to arrive at a final full-spectrum simulated image.

We produced three different planar pupil masks. One corresponds to the full useful aperture of the lens, with an 8-mm pupil diameter. The second mask is an axially-centered pupil with a 1-mm diameter, the size at which diffraction and chromatic effects are comparable. The third mask approximates the U-shaped semi-annular pupil seen in many free-swimming diurnal cephalopods under bright illumination (shallow-water squid and cuttlefish), with a 6-mm inner diameter, a 6.66-nm outer diameter, and a polar angle extent of 180 degrees.

Our MATLAB program uses the wavelength-dependence of the focal length of the *Octopus australis* eye as reported (*22*) by Jagger & Sands. Their laboratory measurements show a sub-percent perturbation in focal length due to residual spherical aberration, but a chromatic fractional shift in focal length, δ*f/f*, of 4.1% between 450 and 700 nm. A second-order polynomial fit to the data in Jagger & Sands yielded δ*f/f* = −5.4676×10^−5^λ^2^ + 0.0794λ − 27.1047, with δ*f/f*=0 at 550 nm.

We modeled a cephalopod lens with a 10-mm diameter d and a focal length of *f*=12 mm at 550 nm, measured from the center of the lens. The spectral range used in the computer model was restricted to 450nm<λ<650nm in order to avoid making an extrapolation from the measured chromatic focal changes reported by Jagger & Sands (*22*). Substantial amounts of illumination (15% of the photons) can lie between 350 and 450 nm, depending on the transmission through the water column. The chromatic focus perturbations are enhanced at short wavelengths, so for our PSF estimates we do extrapolate the Jagger & Sands (*22*) data down to 350 nm, using the expression given above, in conjunction with the attenuated photon spectrum and the opsin response. We used the opsin photon sensitivity curve as a function of wavelength from Chung (*5*).

Our image simulation program used an outermost loop that stepped through a sequence of best-focus wavelengths. This amounts to successively changing the lens-to-retina separation, bringing light of different wavelengths to best-focus at different spacings. For each accommodation value (*i.e*. lens-to-retina separation) we then iterated through illumination wavelengths and computed the appropriate focus offset and blur for that wavelength. The sum, in the wavelength direction, of the blurred hyperspectral image stack produced a 2-D simulation of the test pattern image on the retina, integrated over illumination wavelengths and the opsin response function, for 450<λ<650 nm. We took care to introduce appropriate parity flips of the annular pupil (apparent in Figs. S4-S5) according to the sign of the focal length offset at each wavelength. The wavelength-summed images were normalized so that the intensity value in a resolved test bar was unity.

Representative PSFs for the semi-annular pupil and a point-like white reflector, R(x,y,λ)=δ(x,y), are shown in Figs. S4-S5. The PSF we computed when 400-nm light is brought to a sharp focus (Fig. S4) shows the spike from the point source at 400 nm superimposed on the out-of-focus pupil images from the other wavelengths, from either side of focus. The PSF obtained when the lens is far out-of-focus (Fig. S5) is in effect a radially dispersed image of the point source. These intensity distributions are not well-represented by Gaussian PSFs, and so we used encircled energy (Fig. S7) computations to produce a Gaussian-equivalent Full Width at Half Maximum (FWHM) for these PSFs. The PSF produced by a large circular pupil (Fig. S6) is axisymmetric, and breaks the relationship between radial position and wavelength. The light at the center of the PSF contains all wavelengths, but only from the rays that pass close to the optical axis.

### Contrast Transfer Function (CTF) Analysis

The blurred test pattern images were each analyzed to assess the sharpness of the image, using line profile plots across the images (Fig. 4). We defined a CTF metric that has the merit of being simple and that suppresses aliasing artifacts from the bar pattern and the pupil shape; we computed two times the standard deviation of the pixel values in each image. A crisp image has a bimodal normalized intensity histogram (predominantly 1’s and 0’s) and a high standard deviation. A highly blurred image has an intensity histogram peaked at the mean pixel value, and a low standard deviation. By mapping out this CTF metric vs. lens-to-retina spacing, we can quantitatively assess the extent to which image sharpness can be used to deduce scene spectral content. These results are presented as CTF vs. accommodation plots for various pupil shapes and simulated scene spectral content in Fig. 4.

The animation Movie S3 shows how the contrast of the simulated image depends on focal setting, for the Black-Yellow test pattern shown in Fig. 4.

### Image Quality Budget

We evaluated the various terms in the image quality budget (Table S1) using geometrical or diffractive optics principles, as appropriate. Each Table entry is provided as Gaussian-equivalent FWHM in the focal plane, in units of microns, for the *f*/1.2 spherical lens and a 12-mm focal length, with a radial gradient in index of refraction that compensates for spherical aberration. To convert to an equivalent angular resolution (which is independent of lens diameter for those terms that are in the geometrical optics regime), FWHM∠=2arctan(FWHM/(2FL)).

Adding all the blur contribution in quadrature for the annular pupil case yields an angular resolution of FWHM∠=0.3 degrees. This is in broad agreement with behavioral experiments (*48*) that indicated a dynamic minimum separable angle (MSA) measured for 80mm mantle length animals, in bright broadband light in the cephalopod *Sepia officinalis* of MSA=0.6±0.2 degrees. A determination of cephalopod acuity on *Octopus australis* was performed by Muntz & Gwither (*49*) using static resolution targets. If we take the minimum detectable static contrast for cephalopods to be (*6*) 15% then their results correspond to an angular FWHM of 0.2 degrees, again in basic agreement with our image quality budget estimates, showing that there is no other term greatly compromising the image quality perceived by these organisms.

The geometrical optics approximation (and the diameter-independence of FWHM∠) breaks down if a physical length scale becomes important. There are two limiting cases where that occurs: (1) for sufficiently small (d<1mm) pupils, the wavelength of light becomes important and diffraction dominates the image quality budget, and (2) the fixed photoreceptor size limits angular resolution for lens diameter d<1mm.

Despite the single photoreceptor type, chromatic aberration dominates the image quality budget except for a small, on-axis pupil shape.

#### Photoreceptor size

The typical diameter for the rhabdomes tiled across the cephalopod retina is reported (*11*) to be 5 microns. This sets a limit on spatial sampling in the focal plane. The propagation of light rays across adjacent rhabdomes (*1*) (“rhabdome crosstalk”) would induce additional (and potentially chromatic) image degradation, but studies of cuttlefish retina concluded (*46*, *50*) that their rhabdomes are clad in pigmented sheathing that may suppress this potential source of image degradation. We therefore elected to not include any potential image degradation from rhabdome crosstalk. If rhabdome crosstalk were a significant contributor to image blur, this would not favor the annular pupil shape seen in these animals.

#### Retinal displacement

Cross sectional light micrograph images of cephalopod retinal structure indicate (*50*) an RMS axial displacement of at most a few microns over spatial scales of tens of microns. This translates into a defocus blur of order 1 micron for the full-aperture pupil. The retinal displacement along the optical axis would have to be comparable to the chromatic focus shift (300 microns, Fig. 3) in order to produce image degradation comparable to the chromatic blur. The “retinal bump” in cephalopods could provide spectral information at fixed accommodation if the line of sight is varied (*32*) so as to shift the scene across this perturbation in effective focal length, or if the object of interest’s image on the retina spans the retinal bump.

#### Residual spherical aberration

Although a spherical lens of uniform index of refraction produces pronounced spherical aberration, numerous studies have shown that the lenses of fishes and cephalopods have a radial variation in the index that largely compensates for spherical aberration. Jagger & Sands show (*22*) a typical FWHM from on-axis residual spherical aberration in octopus of less than 5 microns at full aperture. That was for lenses a factor of two smaller than our 12-mm focal length model, so we have scaled this up to 10 microns for the entry in the image quality budget. We note also that this is for full-aperture imaging, and that the annular pupil greatly reduces the radial span of rays in the system. This is therefore a conservative overestimate for the annular pupil geometry, since the residual spherical aberration scales (*51*) as Δh.

#### Chromatic Aberration

The experimental data (*22*) clearly indicate a wavelength-dependent focal shift in the lens of the octopus. We used our quadratic fit to the fractional chromatic focal length shift measured for octopus lenses from Jagger & Sands to perform a numerical computation of the 80% encircled energy radius for a point source at infinity, with best-focus accommodations corresponding to wavelengths between 350 and 650 nm, for the three different pupil geometries we studied. For this computation we were interested in the entire wavelength range of interest, and so we extended the focal length dependence on wavelength down to 350-nm wavelengths. Representative PSFs for accommodation settings that correspond to 400 and 700 nm are shown in Figs. S4-S6._ The encircled energy as a function of distance from the centroid is shown in Fig. S7. We converted from the 80% encircled energy radius, R80, to a Gaussian-equivalent

FWHM=1.3*R80. This produced FWHM Gaussian equivalents of 6, 48 and 61 microns for the small, full and annular pupils, respectively, at the best-focus wavelength of 500 nm, the peak of the opsin curve.

#### Diffraction

The diffraction limit on the focal plane has a spatial FWHM given by FWHM_diff_ = (*f*/)(λ). For our full-pupil with d=8 mm, at the wavelength of peak opsin sensitivity this gives FWHM_diff_ = 1.5*0.5microns = 0.75microns. Stopping down the pupil to a smaller diameter d increases this term by a dimensionless multiplicative factor of (8 mm/d). Our smallest circular pupil diameter of d=1mm produces a diffraction limit of 6 microns FWHM, which is equal to the chromatic aberration term at that small aperture.

#### Other Achromatic Aberrations

In order to constrain the magnitude of aberrations other than those listed above, we turned to the narrowband measurements of Cephalopod PSFs performed (*26*) by Gagnon *et al*. They measured the FWHM produced in collimated 550-nm light (with a 10-nm bandwidth), at full aperture. Their data show a strong correlation between FWHM and *f*-number. For their results on *f*/1.2 lenses, such as the one modeled here, when scaled to a 12-mm focal length they observe a FWHM of 20 microns. We have therefore entered this value in Table S1, under “other achromatic aberrations.” We do not know the ray height dependence of these other aberrations, but unless the blurring induced by these other sources of wavefront error is independent of ray height h, they will remain subdominant for all pupil geometries. These other contributions add of order 10% (when taken in quadrature) to the total FWHM, and therefore the image quality budget is dominated by the chromatic term.

### Consistency with previous behavioral experiments

Table S2 presents a review of cephalopod behavioral experiments investigating color vision, and describes why these are consistent with our proposed mechanism. Before the determination that cephalopods possess a single photoreceptor type, there were numerous experiments demonstrating they had spectral discrimination. These were summarized and dismissed in a 1973 paper (*8*) by Messenger *et al.* with the following rationale:

> “…all the authors are guilty of one or more of three serious errors: failure to take into account the spectral sensitivity curve of the subject, failure to control for the difference in brightness between test objects, and, in the behavioral experiments, inadequate quantification of results, which are presented without conventional statistical analysis.”

We understand that early behavioral experiments demonstrating color vision through training are difficult to validate given other potential cues the animals may use. However, this critique (*8*) of Kühn’s 1950 paper (*20*) is inaccurate and his study is mischaracterized (*8*) in their Table 1 as purely a training experiment. A reading of Kühn (*20*) shows that while he did perform extensive training experiments indicating spectral discrimination in *O. vulgaris*, he also clearly demonstrated the differential responses of cuttlefish chromatophores to differentially colored textured backgrounds. A translation (*52*) of the relevant section of Kühn (*20*) states:

> “Both the statistical recordings of the observed color, that are shown in figures and tables, and the immediate observations of the chromatophore changes show, that colored environments provoke different reactions than non-colored luminances. The differences in gray value on the grayscales get answered by a differential expansion of the black chromatophores. Their differential expansion also shows a different ‘luminance-value’ in the colored environment; yet added to that is a counteracted behavior of the colored chromatophores: if the color of the substrate is of short wavelength (blue, green), the colored chromatophores, the orange colored ones extremely, become small, if the color is of long wavelength (yellow, red), the colored chromatophores become extended, the more, the longer the wavelength of the light that they reflect. These reactions prove a ‘Sense of color’ of the cuttlefish, i.e. the capability to discriminate certain wavelength qualitatively from luminances of mixed light that appears colorless.”

In our proposed mechanism cephalopods cannot gain spectral information from either a flatfield background or an edge between two abutting colors of comparable intensity (Fig. 3). This would explain why optomotor assays and camouflage experiments using abutting colored substrates (*6*, *8*, *10*) fail to elicit a response different from a flat-field background. Similarly, experiments (*9*) with monochromatic light projected onto a large uniform reflector or training experiments (*7*–*8*) with rapidly vibrating colored cues would defeat a determination of chromatic defocus. Cephalopods fail tests for color vision in these special cases that would defeat our proposed mechanism yet succeed in scenarios where the animals are given focusing aids to determine chromatically-induced defocus.

**Figure S1.**
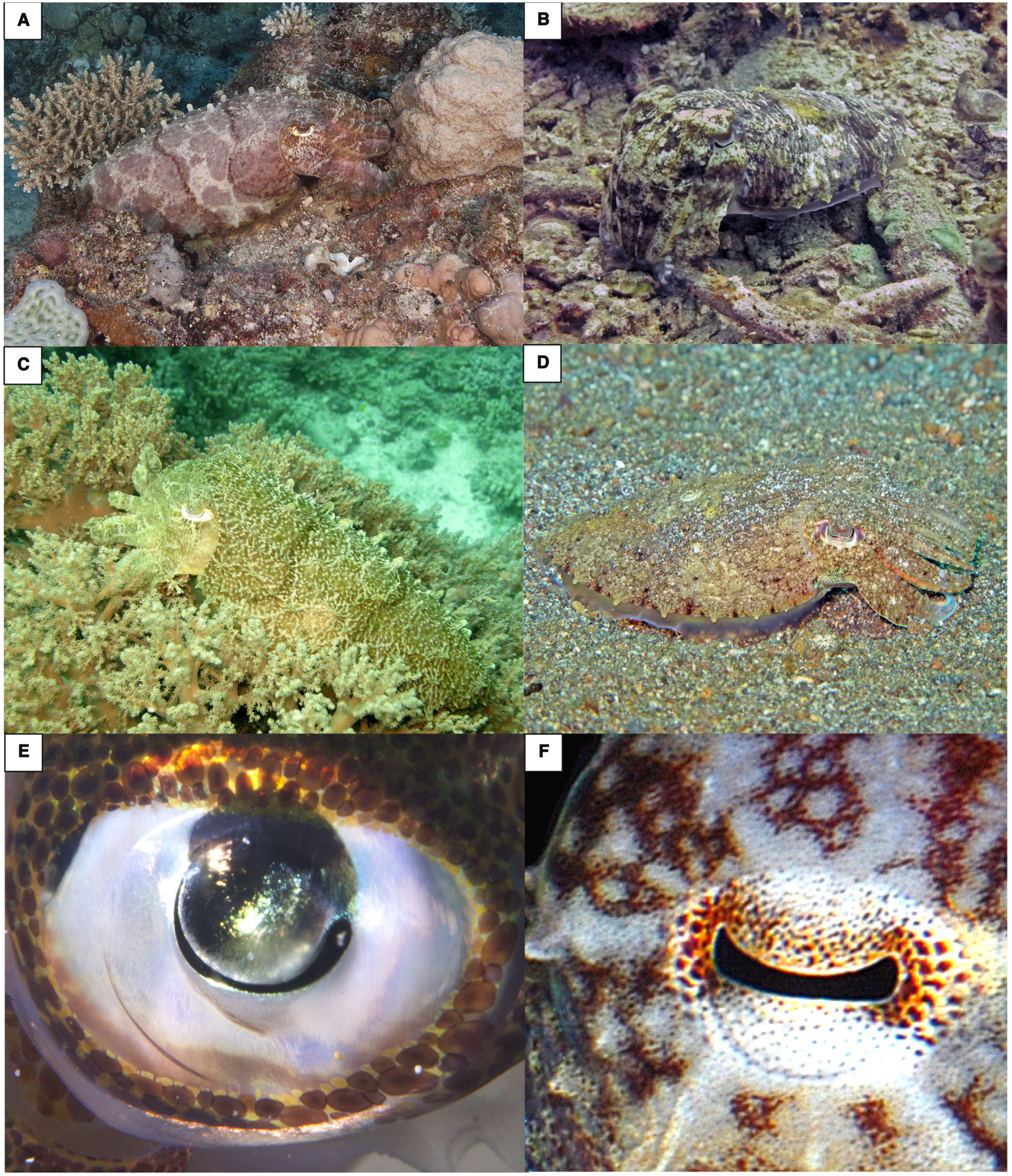
Color matching and pupil shape in cephalopods. (**A-D**) The coral reef broadclub cuttlefish *Sepia latimanus* lives in one of the most chromatically complex ocean environments, and accurately color-matches natural environments to camouflage. (**E**) The pupil in shallow-water squids (such as this *Sepioteuthis*) maximizes off-axis light when imaging in the horizontal plane. (**F**) Shallow-water octopus species such as this *O. rubescens* maximize off-axis light when imaging the bottom. More examples of color matching in cephalopods are found in Movie S1.

**Figure S2.**
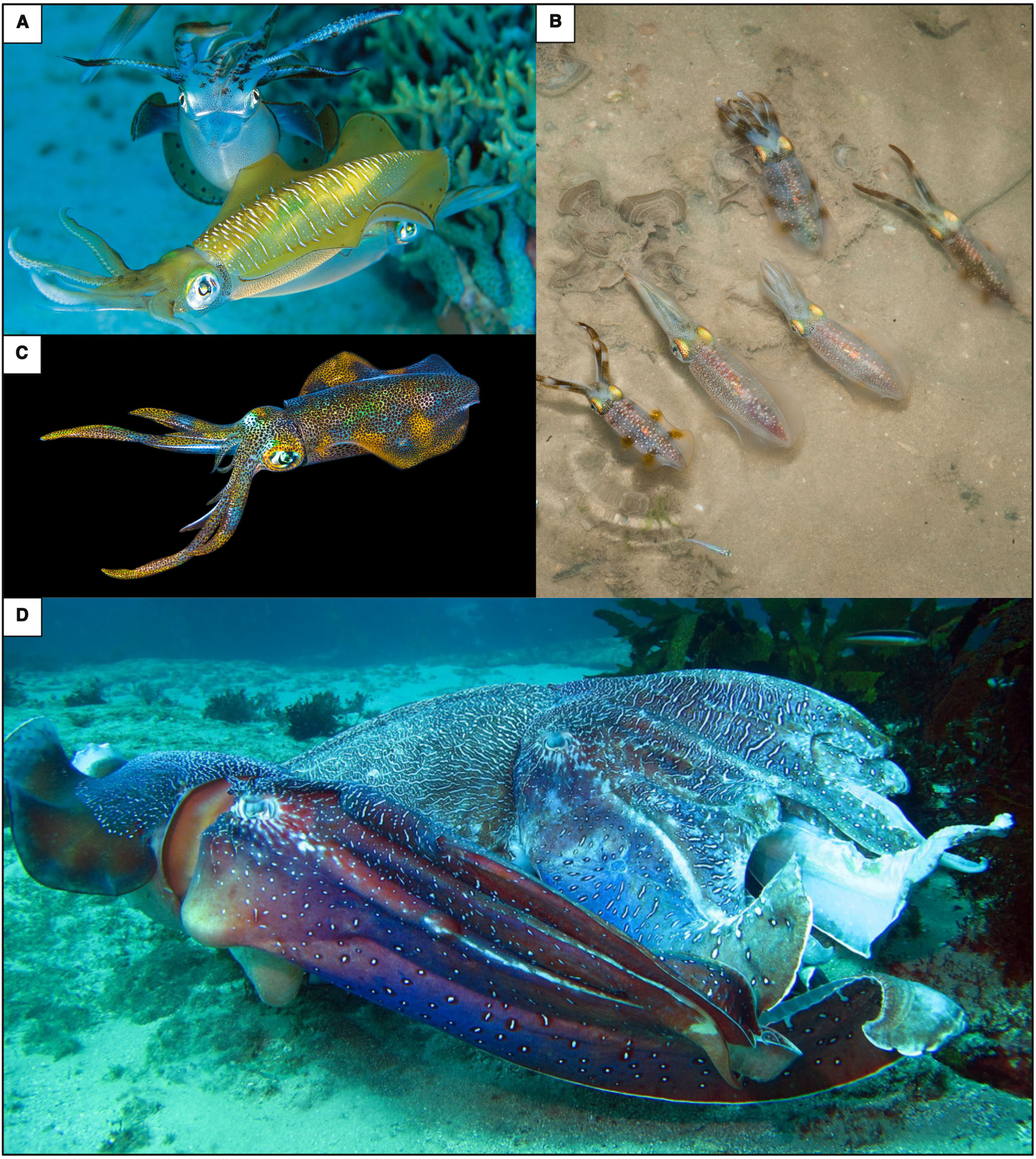
Examples of highly colorful signaling in cephalopods. (**A-C**) The bigfin reef squid *Sepioteuthis lessoniana* vividly changes color while signaling to members of its own species. (**D**) The Australian giant cuttlefish *Sepia apama* also uses a colorful display and the fine network of black lines allows for an easy determination of chromatically induced defocus and thus chromatic information. During displays the cuttlefish pupil is typically maximally contracted as a semi-annulus, suppressing other sources of image degradation while maximizing chromatically-induced blurring. More examples are found in Movie S2.

**Figure S3.**
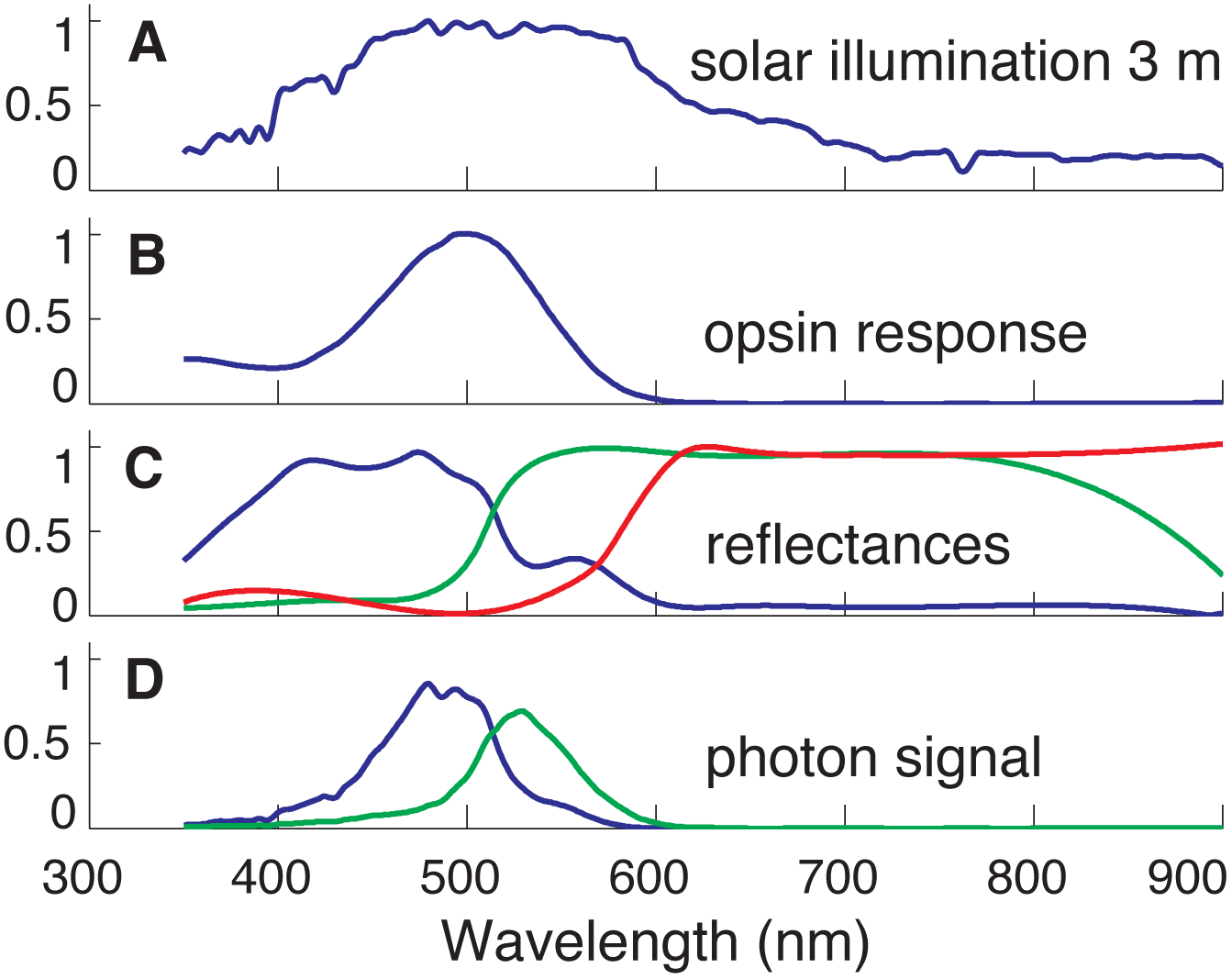
Spectra used in simulations showing the wavelength dependence (in nm) of multiple quantities of interest. (**A**) Depth-attenuated solar photon spectrum. (**B**) The sole opsin’s relative photon sensitivity response function. (**C**) The three fish reflectance spectra we used. (**D**) Detected photon flux for the two bluest reflectance spectra that were used for the test patterns shown in Fig. 4. All plotted quantities are dimensionless and normalized.

**Figure S4.**
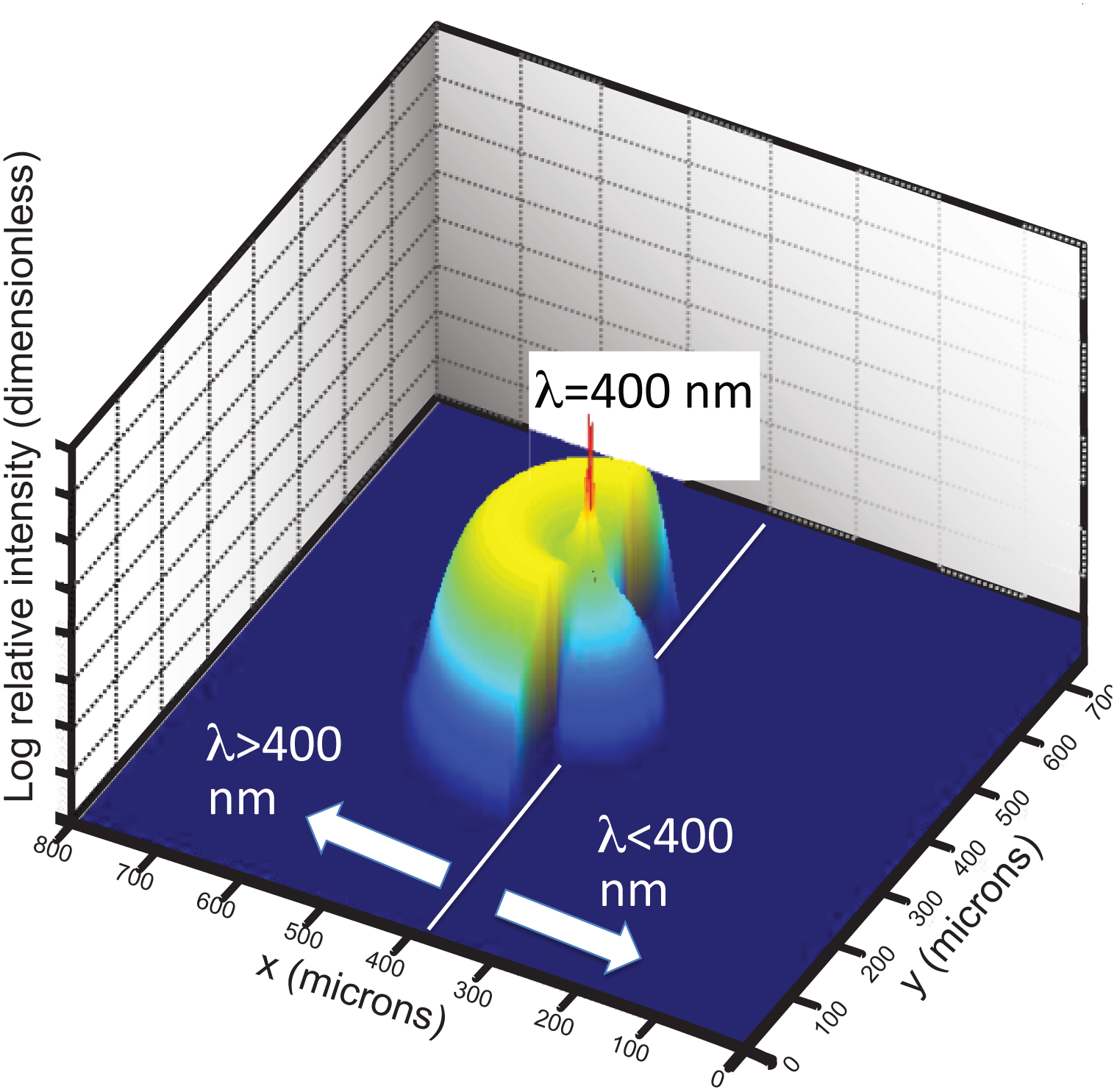
Chromatic PSF for 400-nm best-focus, semi-annular pupil. This figure shows the computed photon intensity on the retinal surface for a point source at infinity, with a solar spectrum filtered to a 3-m depth, convolved with the opsin sensitivity function. The spatial units are microns. This is for the half-annular pupil geometry, so that light with λ>400 nm has yet to come to a focus, while light with λ<400 nm has already passed through its position of best-focus. The spike at the center corresponds to the best-focused photons with λ=400 nm, which have the highest surface brightness.

**Figure S5.**
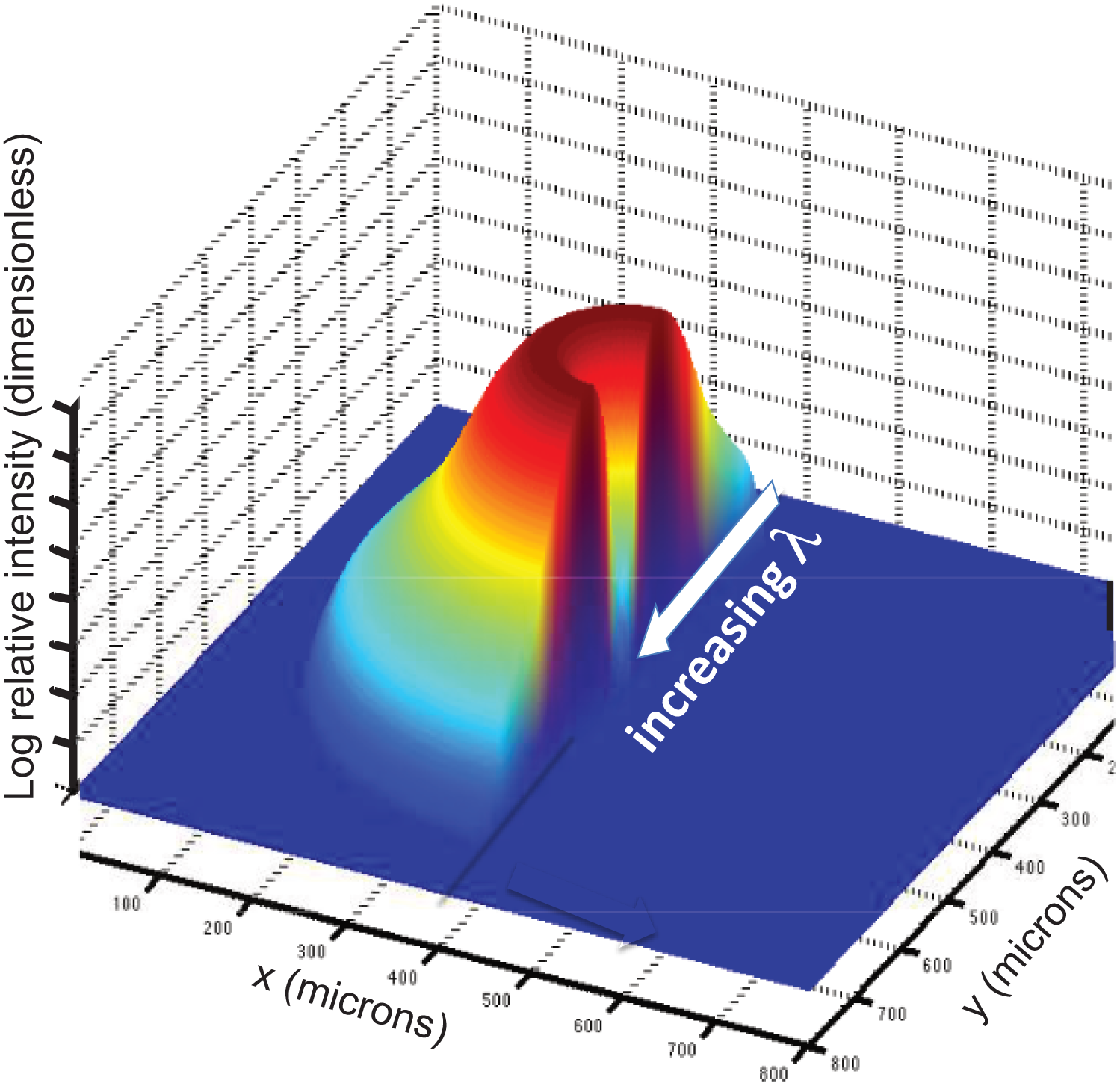
Chromatic PSF for 700-nm best-focus, semi-annular pupil. This is computed for the same half-annular pupil as the previous figure, but the PSF has been rotated about the optical axis by 180 degrees, for clarity. All the rays have passed through their position of best-focus, and so there is a monotonic relationship between wavelength and distance from the axis of symmetry, as indicated. The radial plot of photon surface brightness amounts to a dispersed spectrum of the incident light, with the redder light being closest to being in focus.

**Figure S6.**
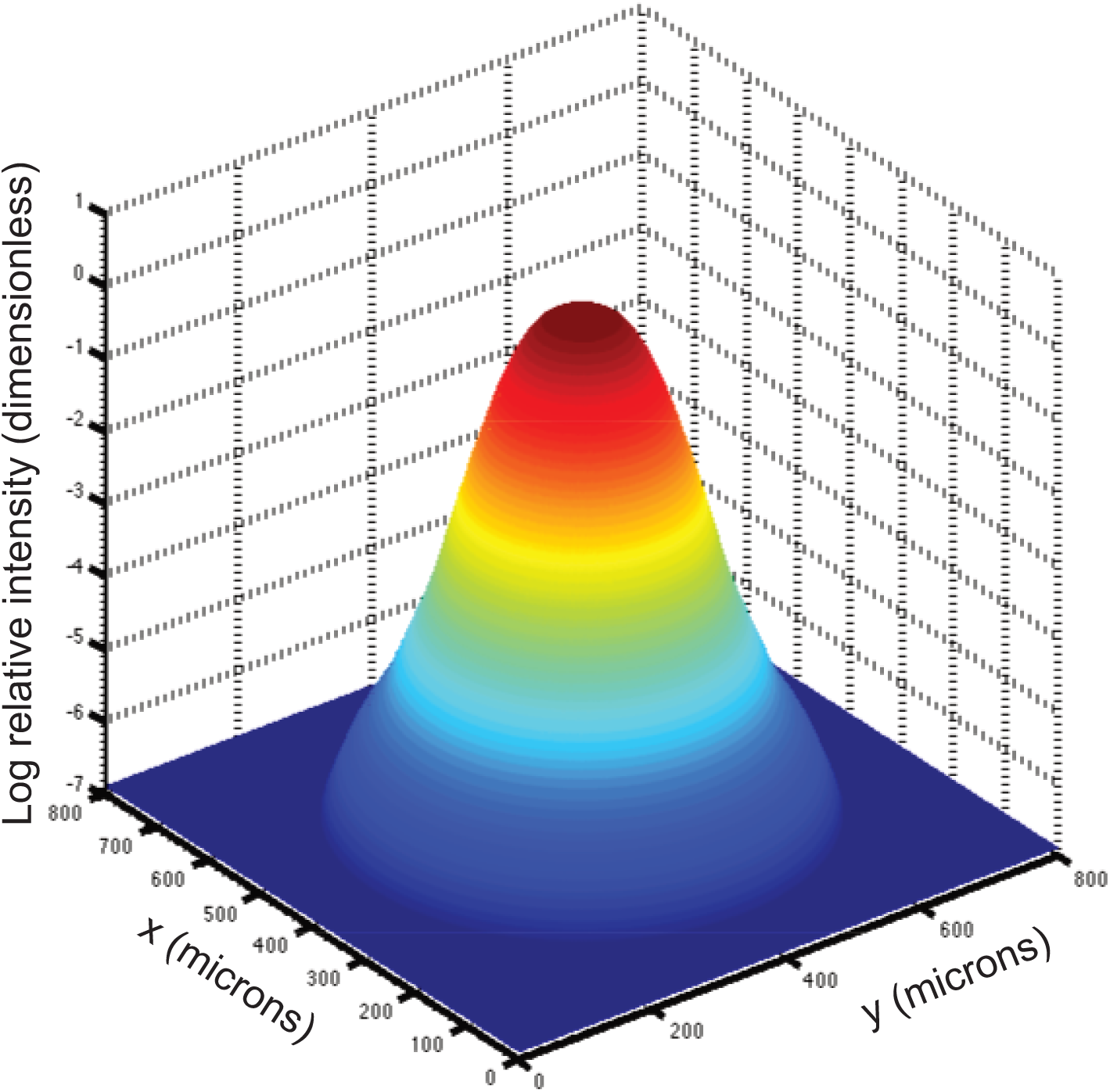
Chromatic PSF for 700-nm best-focus, full pupil. As distinct from semi-annular case, the center of the PSF is filled in with polychromatic light that passes through the center of the pupil and suffers minimal chromatic blurring, while the outer edges of the PSF are illuminated exclusively by the photons of the shortest wavelength. There is therefore a radial gradient in spectral purity and a complex relationship between position and spectral content.

**Figure S7.**
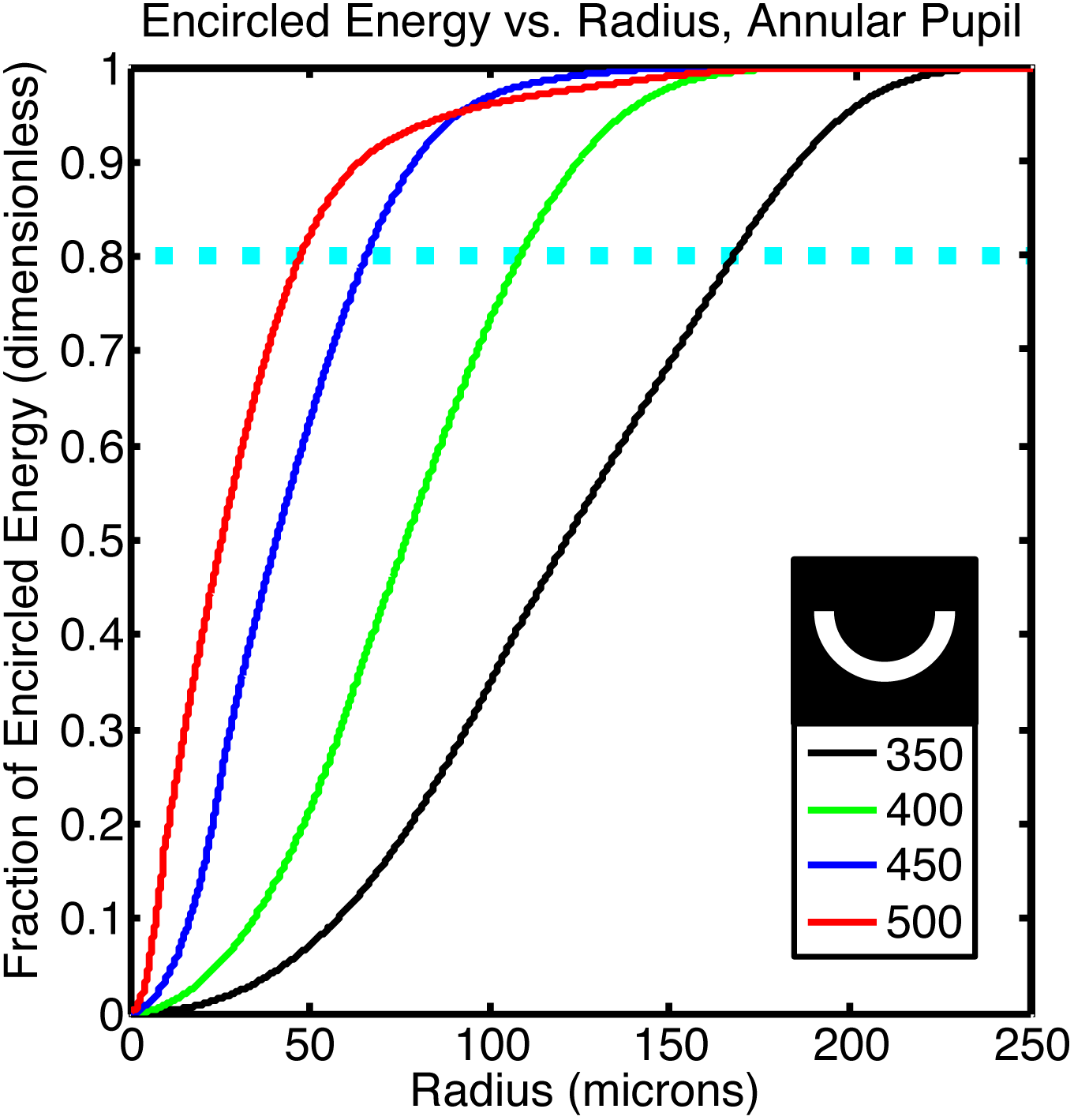
Encircled energy vs. radius for various accommodation settings. This result is for the semi-annular pupil at different accommodation settings indicated as the best-focus wavelength in nm. The plot shows the integrated enclosed energy within the PSF for the semi-annular pupil geometry. This was computed for a white reflector illuminated by the depth-attenuated solar photon spectrum, for 350<λ<650 nm. The red curve yields an 80% encircled energy radius of 47 microns, which corresponds to a Gaussian-PSF-equivalent FWHM of 61 microns, at the accommodation setting of sharpest focus, which corresponds to a best-focused wavelength of 500 nm.

**Table S1.** Cephalopod retinal image quality budget. The columns present the various aberration phenomena, the resulting Gaussian-PSF equivalent FWHM, and the dependencies on ray height h and on wavelength λ. Chromatic aberration is the dominant contribution to image blurring, down to 1-mm pupil diameters. The quadrature sum of the various contributions for the semi-annular pupil is 65 microns, and is dominated by chromatic blurring. These values correspond to an *f*/1.2 lens with a 12-mm diameter.

| Term | FWHM ($\mu\text{m}$) | $h$-dependence | λ-dependence |
| --- | --- | --- | --- |
| Photoreceptor size | 5 | none | none |
| Retinal surface displacement | 1 | $\propto h$ | none |
| Crosstalk between photoreceptors | Neglected<br>(see text) | $\propto h$ | unknown |
| Residual Spherical aberration | 10 at full<br>aperture | $\propto \Delta h$ for<br>annular pupil | none |
| Diffraction with $d = 1$ mm, on-axis<br>pupil | 6 | $\propto h^{-1}$ | $\propto \lambda$ |
| Diffraction with $d = 8$ mm, on-axis<br>pupil | 0.75 | $\propto h^{-1}$ | $\propto \lambda$ |
| Other achromatic aberrations | 20 at full<br>aperture | Unknown,<br>likely $\propto h$ | none |
| Chromatic Aberration with $d = 1$<br>mm, on-axis pupil | 6 | $\propto h$ | none |
| Chromatic Aberration with $d = 8$<br>mm, on-axis pupil | 48 | $\propto h$ | none |
| Chromatic Aberration with semi-annular pupil | 61 | $\propto h$ | none |

**Table S2.** Prior behavioral Cephalopod experiments testing for color vision. This Table provides a summary of how prior laboratory cephalopod behavior and vision experiments compare to the chromatic aberration model proposed here.

| Year | Author | Experiment Description | Consistent with Model? |
| --- | --- | --- | --- |
| 1950 | Kühn (20) | a. Test of camouflage response on textured colored backgrounds vs. greyscale | Yes, Kühn concludes a wavelength sensing capability. Both of his experimental designs (**a-b**) allow for the determination of chromatically-induced defocus. |
|  |  | b. Training experiment on *Octopus vulgaris* using stationary targets | Yes (see above). |
| 1973 | Messenger, Wilson & Hedge (8) | c. Training experiments on octopus under fluorescent lighting using colored rods vibrated at 3Hz | Yes, rapidly vibrating color cues would make color assessment via chromatic defocus impossible. |
|  |  | d. Nystagmus response in octopus to alternating colored stripes | Yes, under our model they are unable to judge coloration absent fine scale structure. Adjacent colors are not resolvable. |
| 1975 | Roffe (9) | e. Training experiment on octopus with unfocused monochromatic light projected onto a white screen without focusing cues | Yes, light projected onto uniform disk would not allow for determination of chromatic defocus. |
| 1977 | Messenger (7) | f. Training experiments on octopus with rectangles of colored cues vibrated at 3Hz | Yes, rapid vibration makes color assessment via chromatic defocus impossible. |
| 1996 | Marshall & Messenger (10) | g. Camouflage assay using two adjacent colors of artificial fine gravel | Yes, under our model adjacent colors are not resolvable. |
| 2006 | Mäthger, Barbosa, Miner & Hanlon (6) | h. Nystagmus tracking response in *Sepia* using alternating adjacent colored bars rotated around the head of the animal | Yes, both lines of evidence (**h-i**) use adjacent colors without fine scale structure; this defeats a spectral discrimination model using chromatically dependent defocus as seen in Figure 3. |
|  |  | i. Camouflage assay using two adjacent colors in a uniform checkerboard | Yes (see above). |

**Movie S1. *Octopus vulgaris* demonstrating dynamic background color matching in the wild.** The first segment of this clip shows *O. vulgaris* in shallow water camouflaged against a green clump of algae. In the second segment, the same species rapidly changes to a matching color when faced with a reddish/brown natural background. Used by permission of Dr. R. Hanlon.

**Movie S2. Cuttlefish *Sepia latimanus* demonstrating chromatic signaling to conspecifics.** This movie shows how some shallow-water tropical cuttlefish break camouflage while signaling to conspecifics, in contrast to the same species in camouflage in Fig. S1A-D. While they could easily produce the same luminance function in black and white with less chromatic contrast, as is done in many other *Sepia* species, some shallow-water cuttlefish and squid signal to each other with highly chromatic displays that are conspicuous for their predators with color vision. The chromatic signal is interspersed with black lines, allowing for a determination of chromatically-induced defocus. The colorful mantle signals in the leftmost animal at 0:30 show bright gold and purple separated by a black line, allowing the sign of chromatic defocus to be resolved for each color (while if the two colors were directly adjacent this would be impossible). Note that this movie was filmed in the natural light environment of these organisms, without any artificial lights or filters. This shows that in the natural lighting environment these signals appear “colorful” to vertebrate predators with color vision through multiple photoreceptor channels. Used by permission of J. Aguilera.

**Movie S3. Contrast dependence on focal setting.** This animation shows the simulated results for an annular pupil and the black-yellow test pattern shown in Fig. 4. Notice how the contrast is maximized at the focal setting that brings light at the detected spectral peak into best focus. The upper panel shows the image formed on the retina and the lower panel is a line cut that plots intensity vs. position.

